## Supplementary figures and images for "Fat body-derived cytokine Upd2 controls disciplined migration of tracheal stem cells in *Drosophila*"

### Figure 1-figure supplement 1

##### Figure 1-figure supplement 1

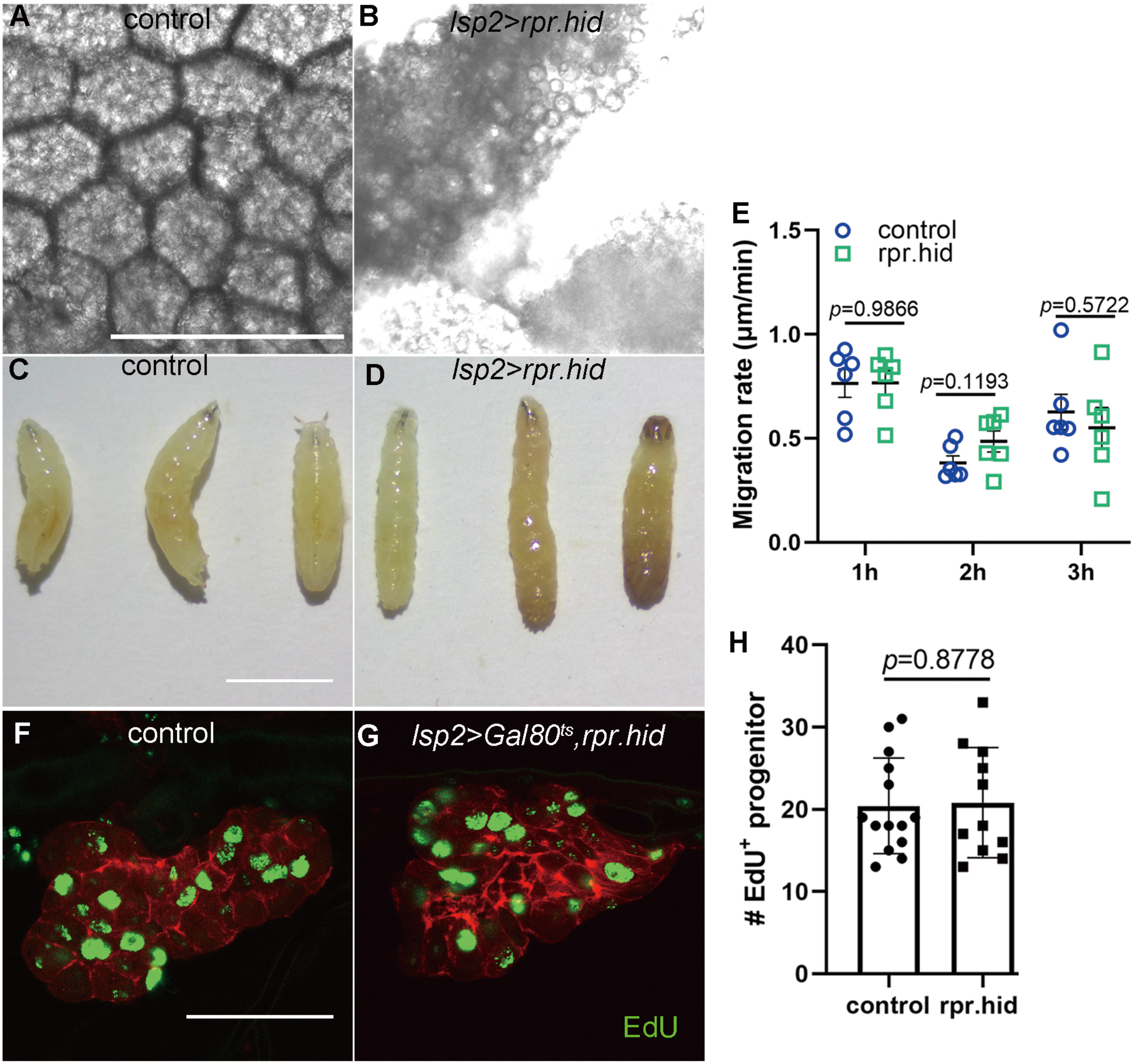

### Figure 2-figure supplement 1

**Figure 2-figure supplement 1**

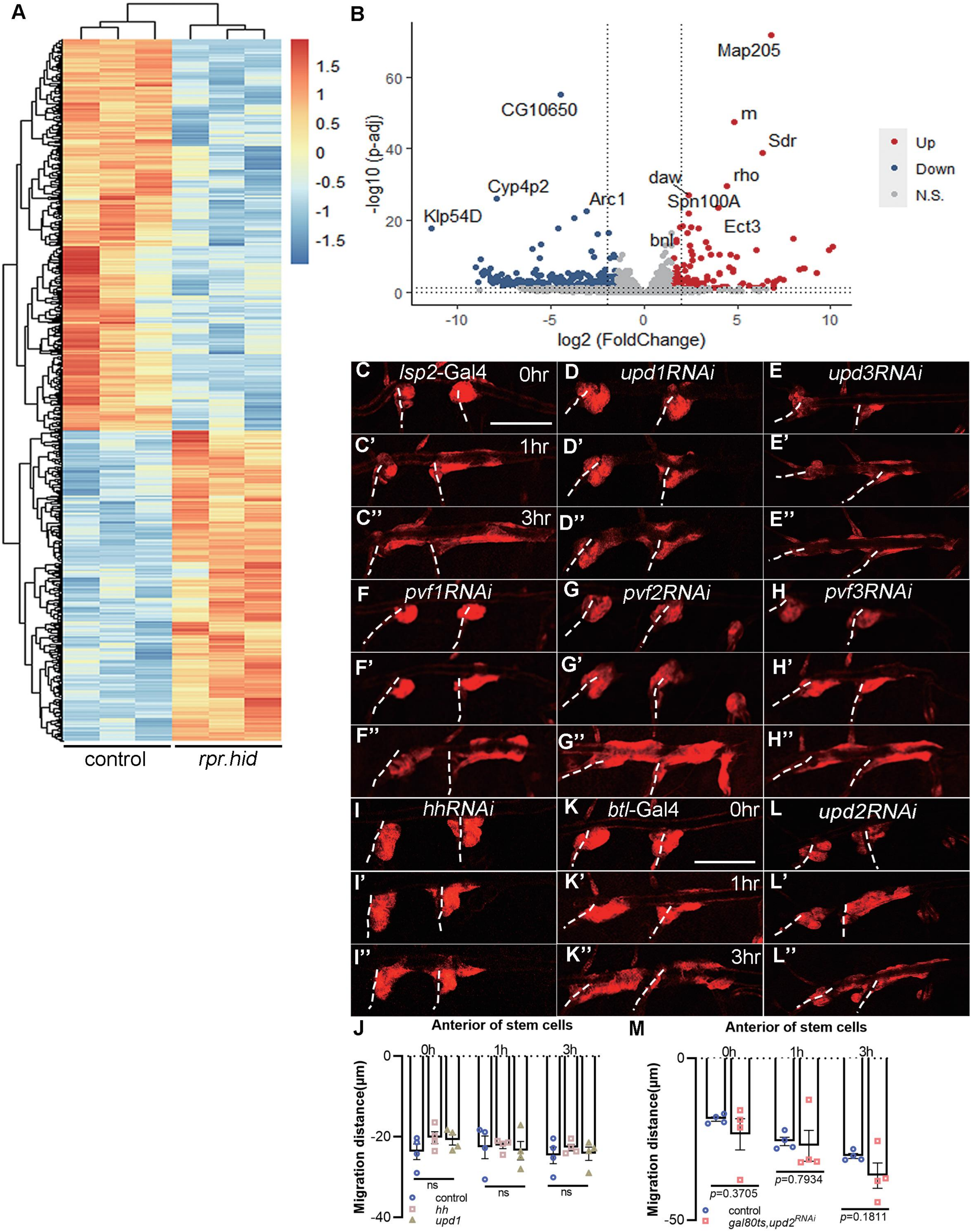

### Figure 2-figure supplement 2

#### Figure 2-figure supplement 2

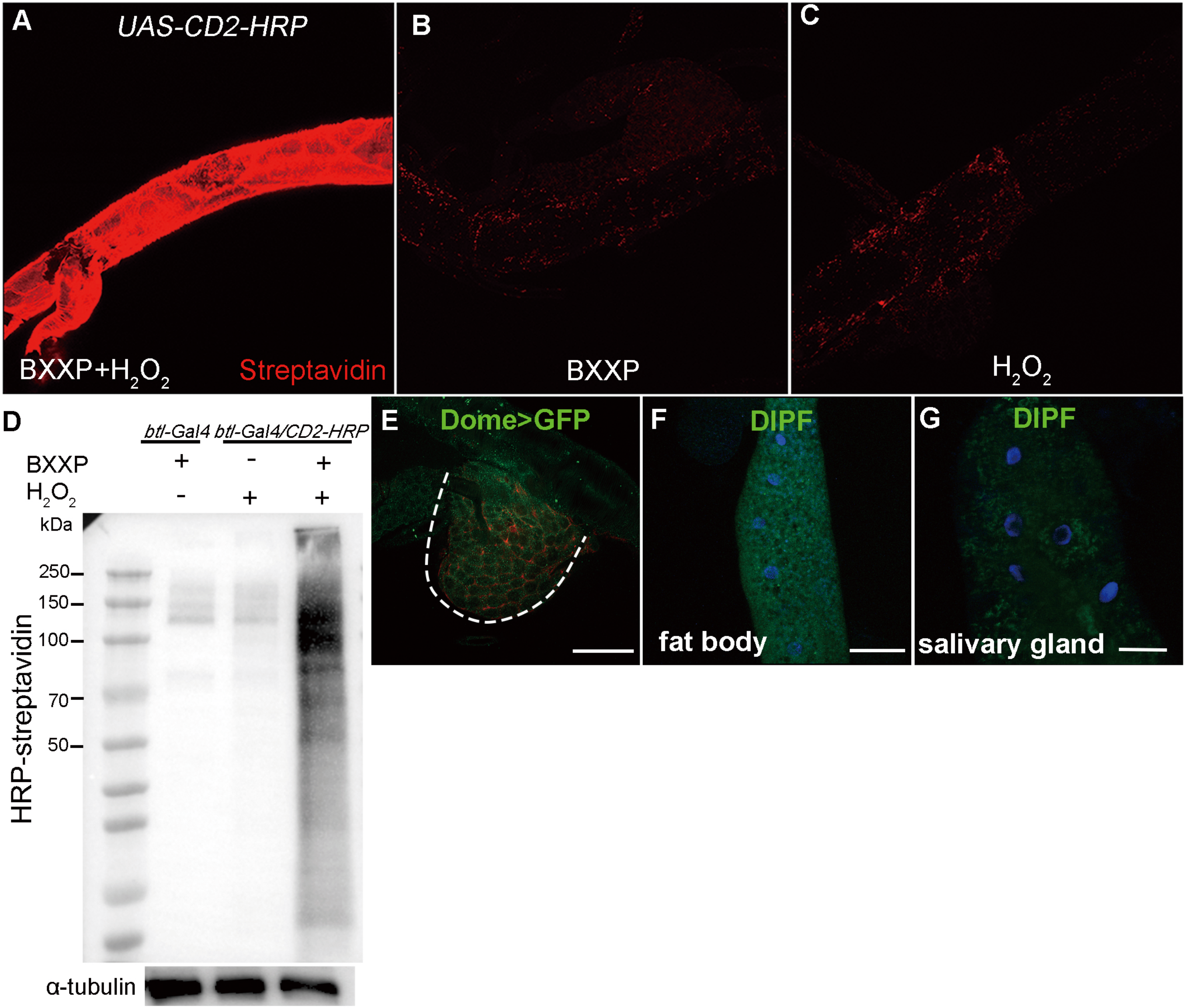

### Figure 3-figure supplement 1

#### Figure 3-figure supplement 1

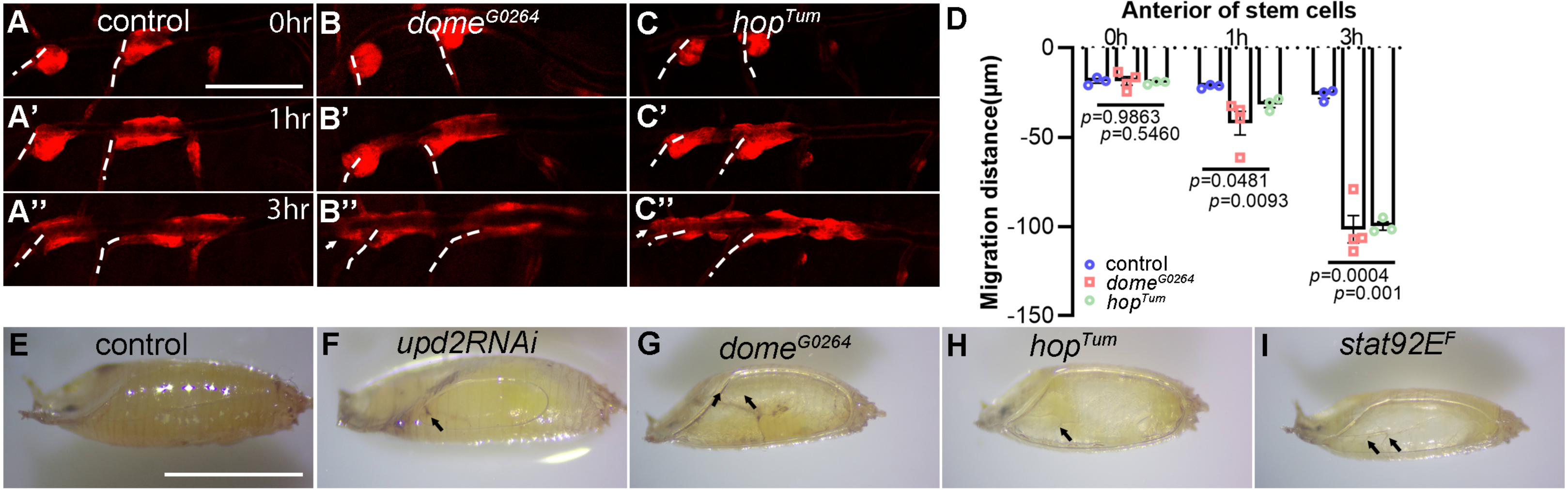

### Figure 3-figure supplement 2

##### Figure 3-figure supplement 2

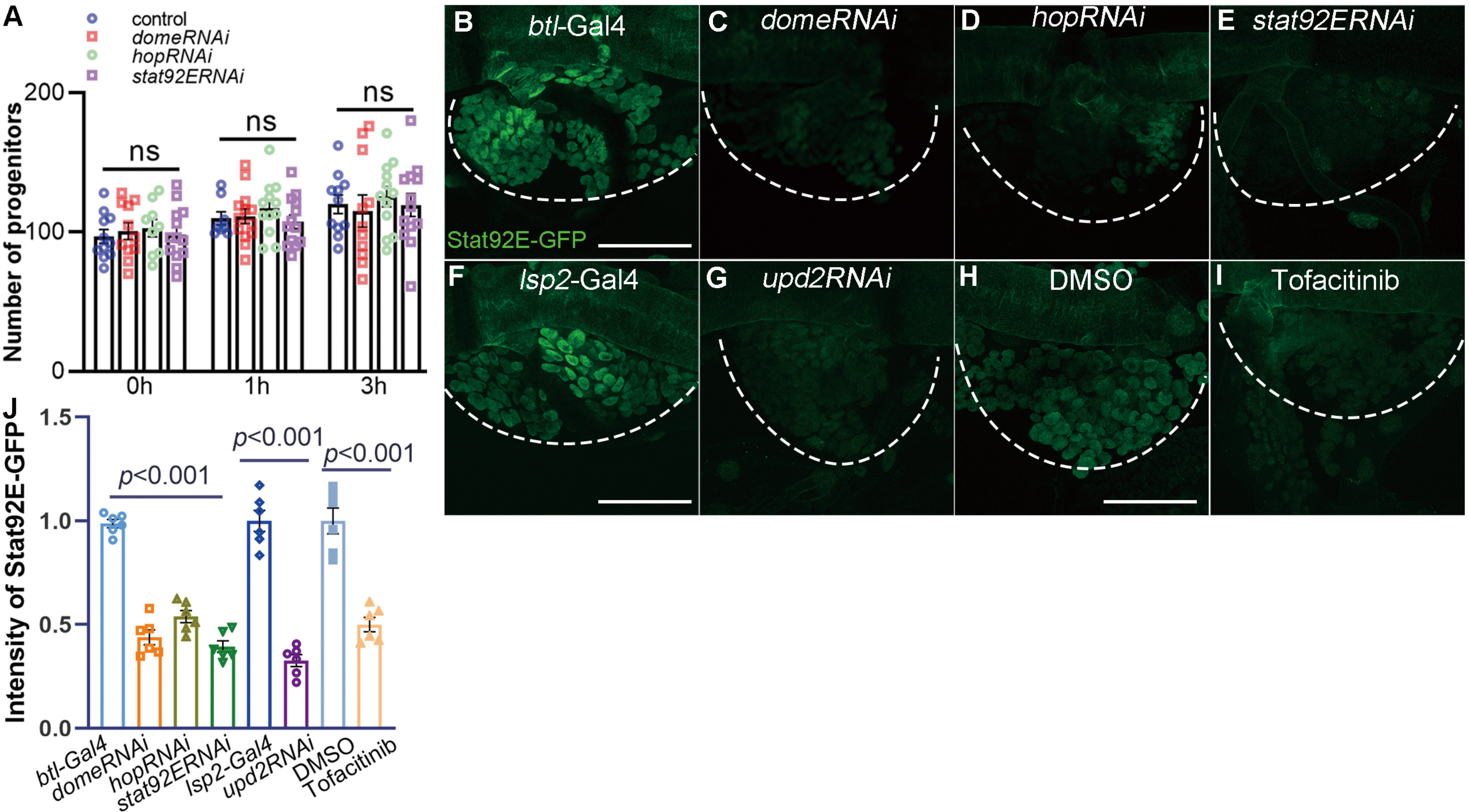

### Figure 4-figure supplement 1

Figure 4-figure supplement 1

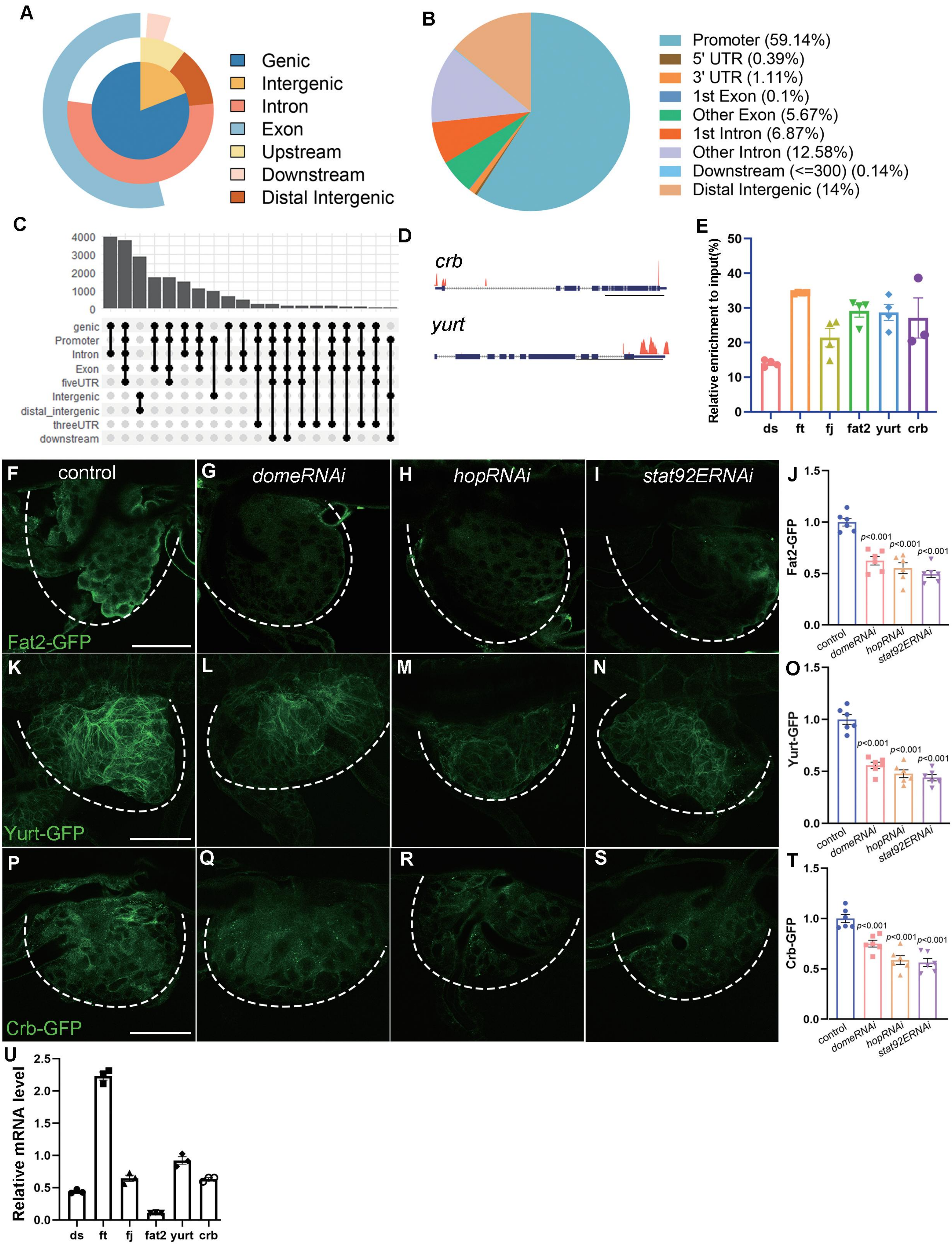

### Figure 4-figure supplement 2

Figure 4-figure supplement 2

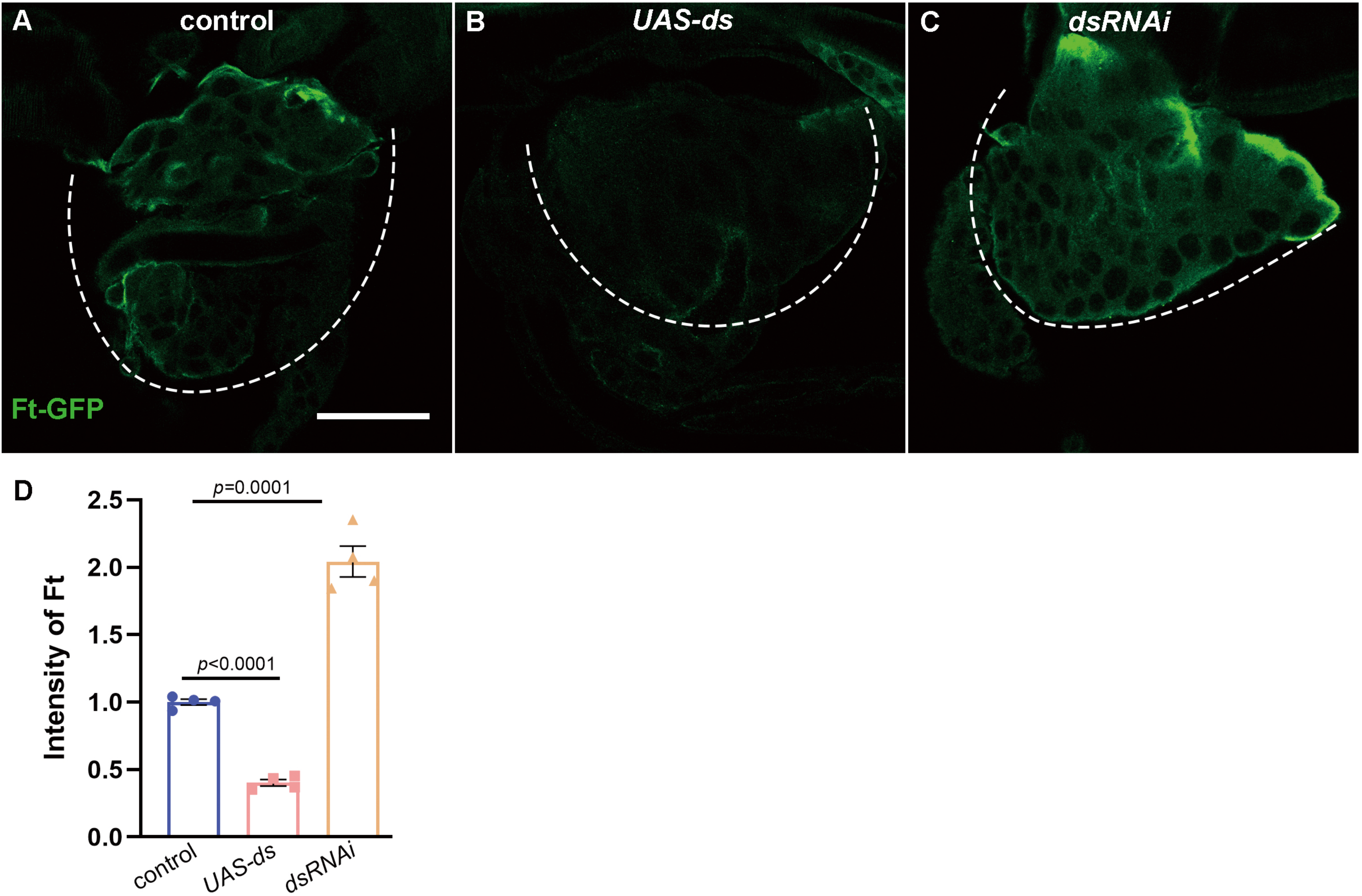

### Figure 5-figure supplement 1

#### Figure 5-figure supplement 1

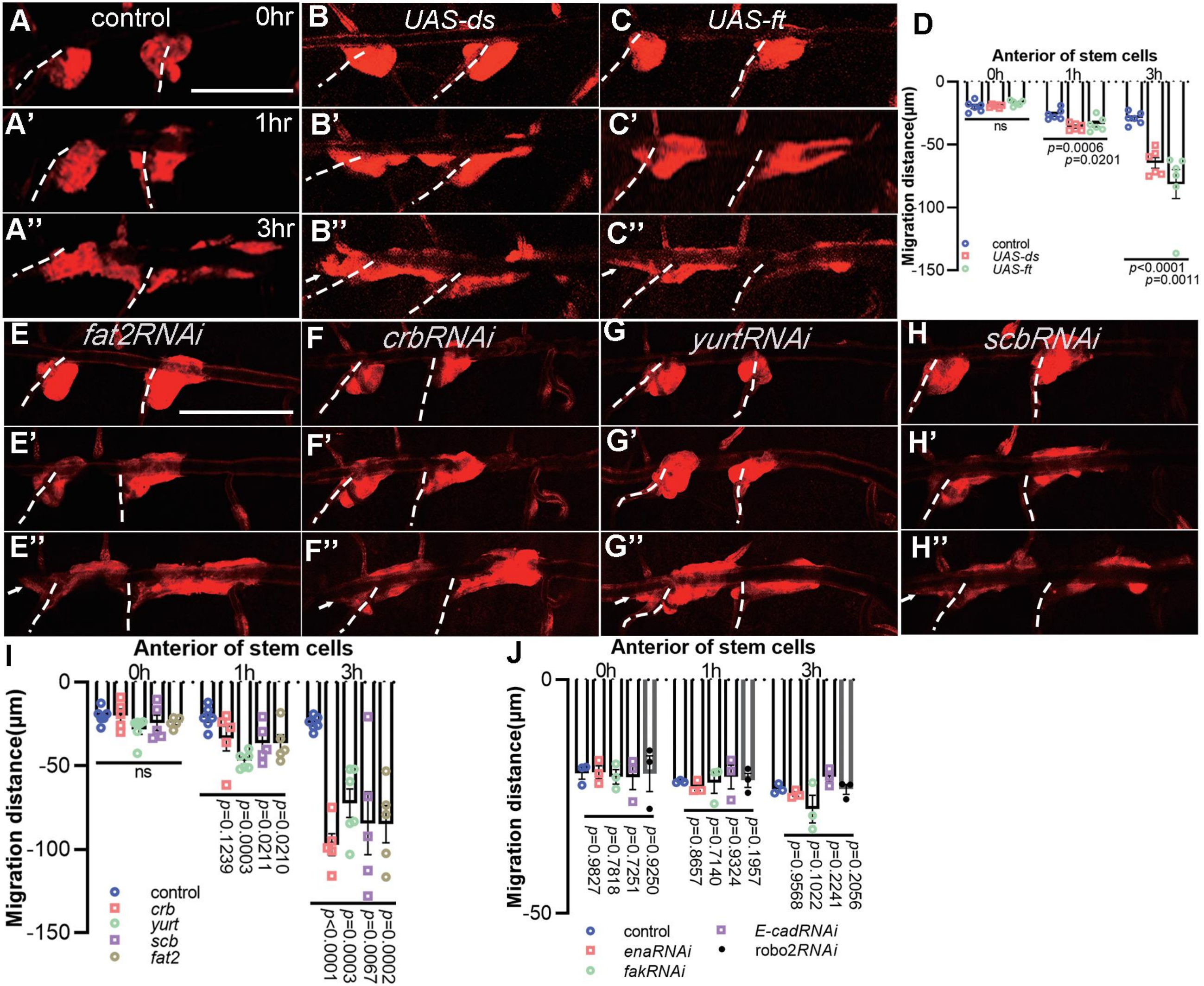

### Figure 5-figure supplement 2

**Figure 5-figure supplement 2**

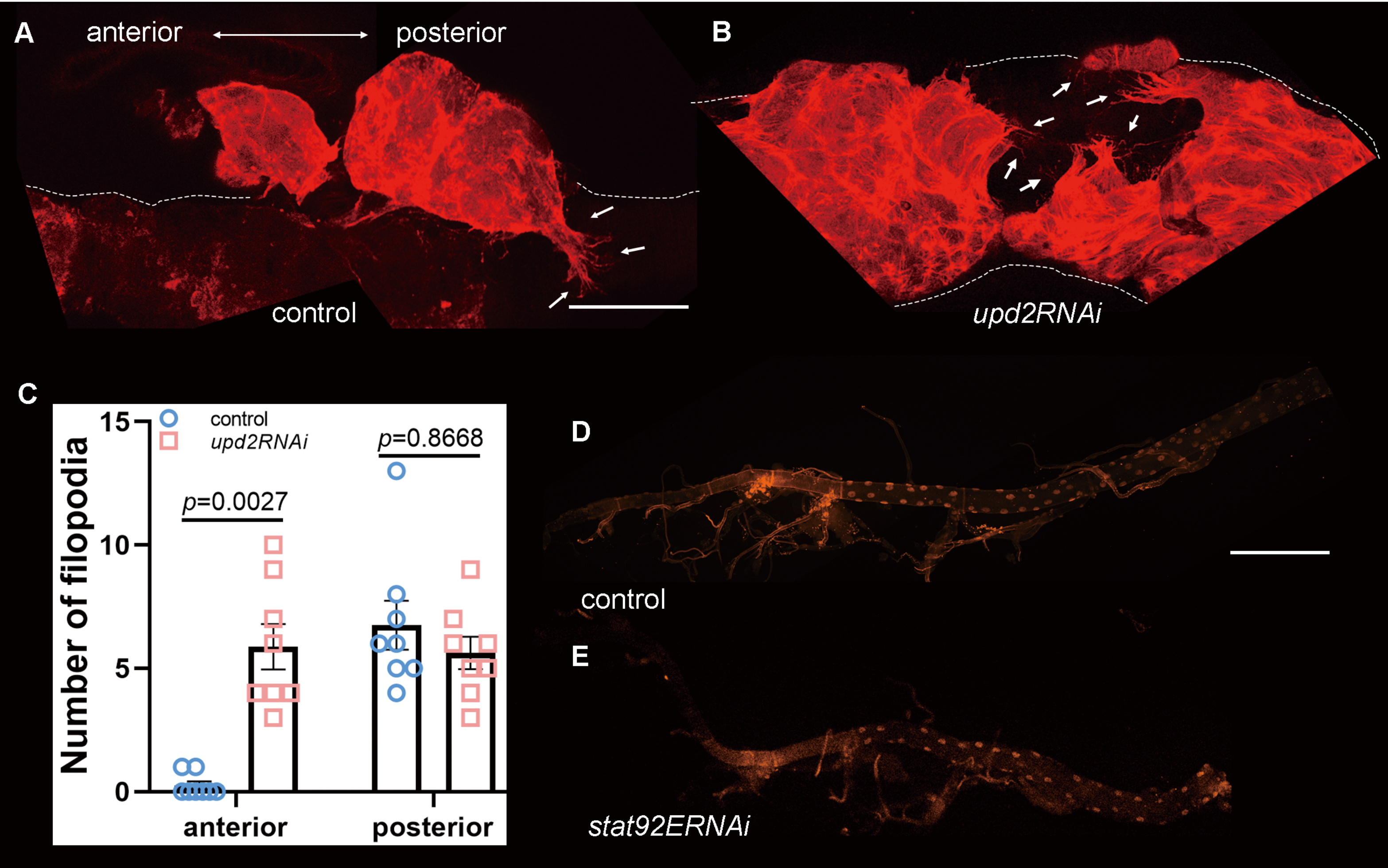

### Figure 5-figure supplement 3

Figure 5-figure supplement 3

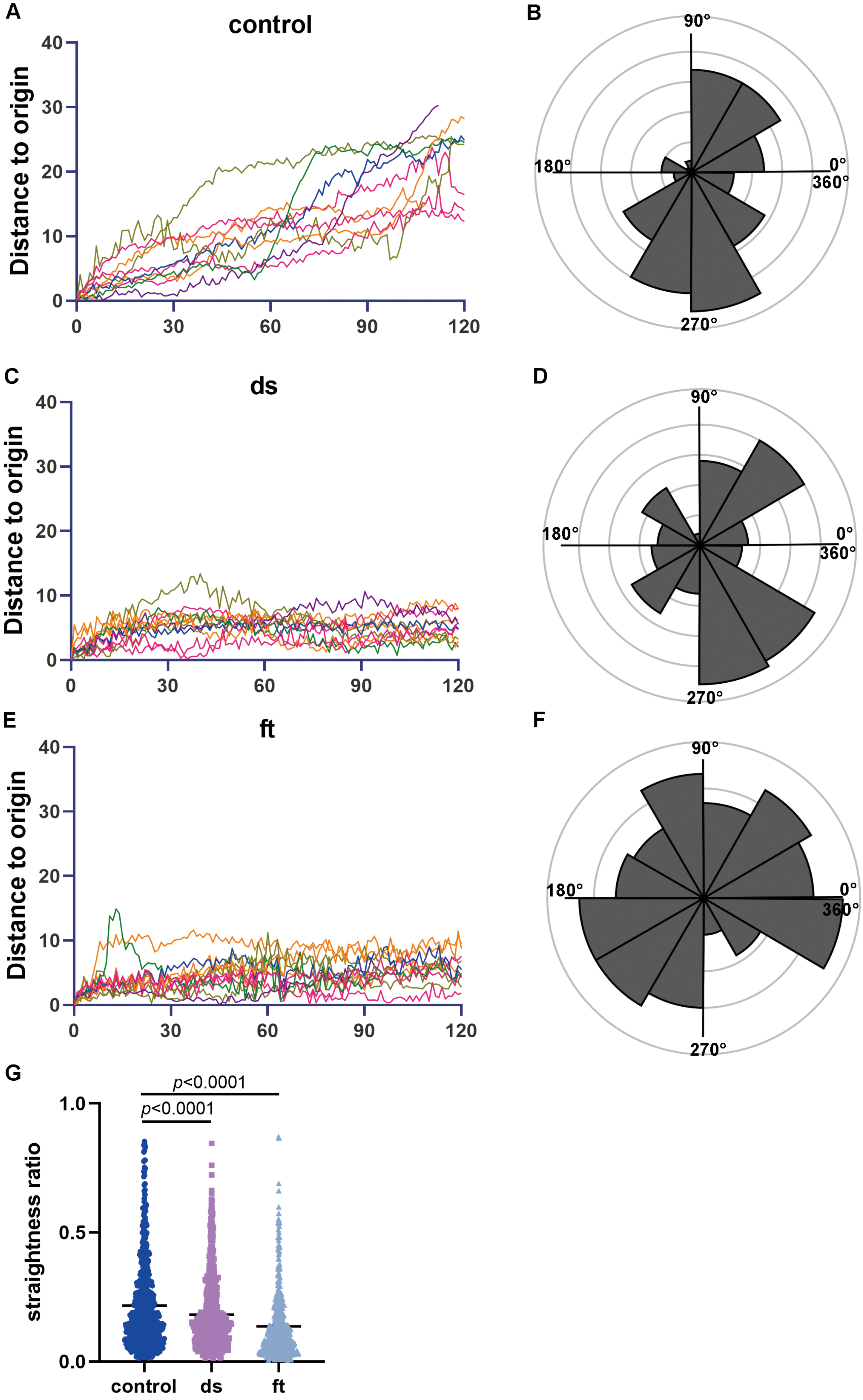

### Figure 6-figure supplement 1

#### Figure 6-figure supplement 1

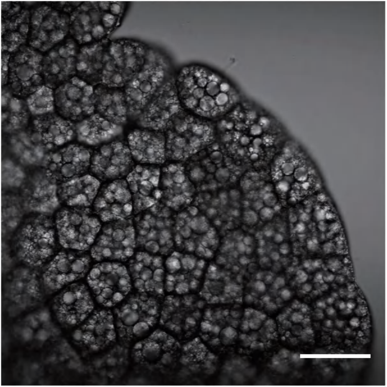

### Figure 7-figure supplement 1

#### Figure 7-figure supplement 1

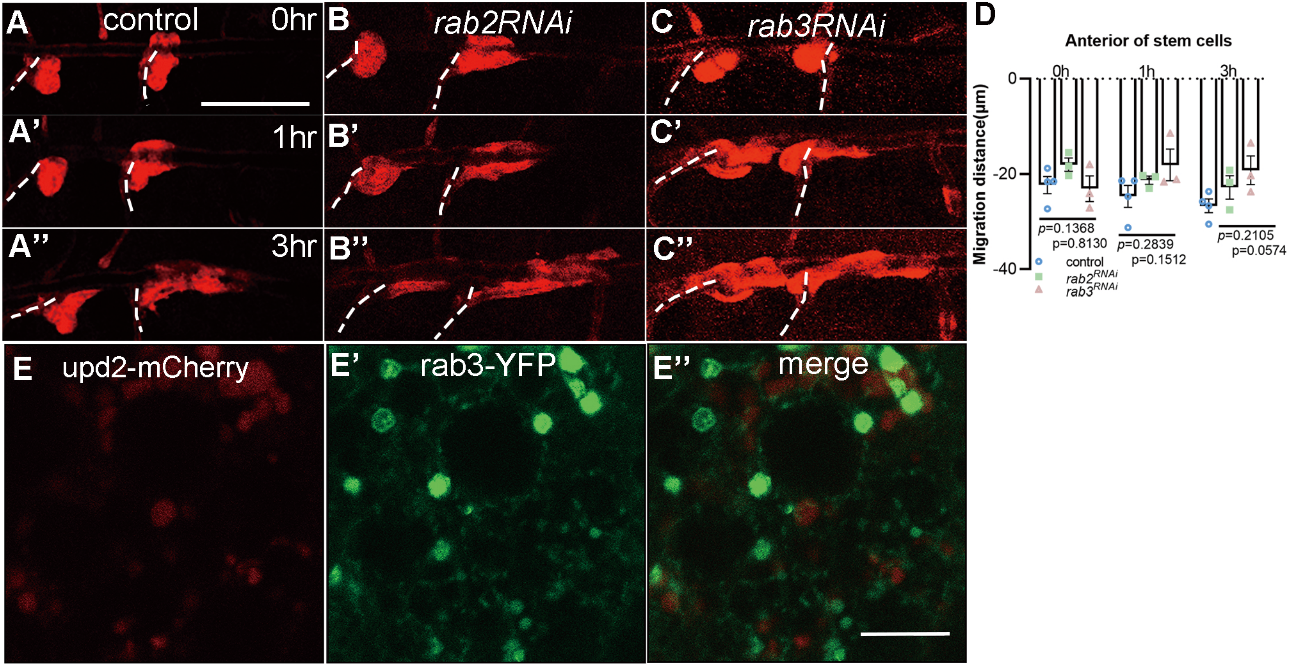

### Figure 8-figure supplement 1

#### Figure 8-figure supplement 1

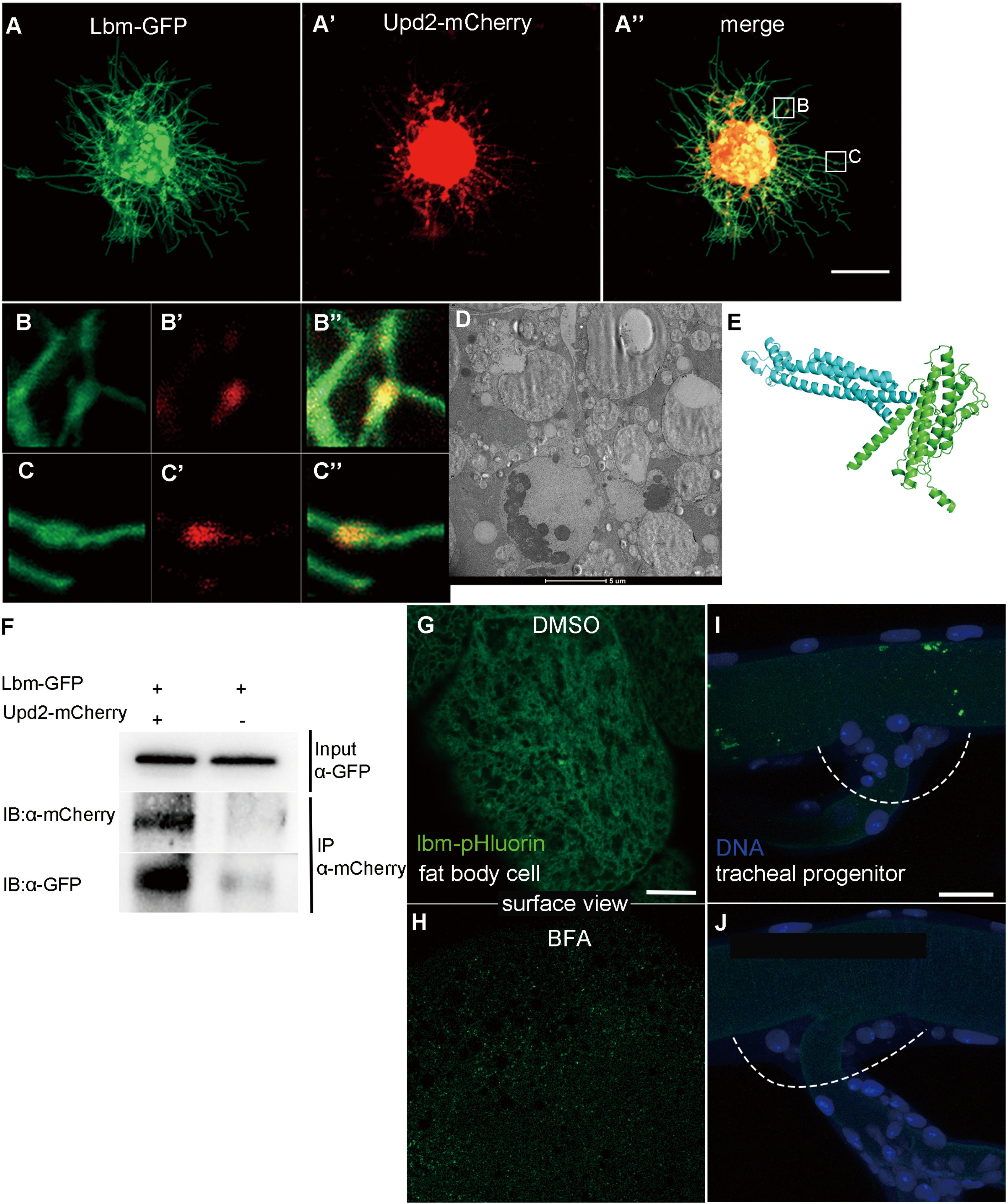
